# Power laws govern the abundance distribution of birds by rank

**DOI:** 10.1101/2021.12.06.471443

**Authors:** Sergio Da Silva, Raul Matsushita

**Affiliations:** Department of Economics, Federal University of Santa Catarina, 88049-970, Florianopolis S.C., Brazil; Graduate Program in Economics, Federal University of Santa Catarina, 88049-970, Florianopolis S.C., Brazil; Graduate Program in Economics, Federal University of Espirito Santo, 29075-910, Vitoria E.S., Brazil; Department of Statistics, University of Brasilia, 70910-900, Brasilia, D.F., Brazil; Graduate Program in Statistics, University of Brasilia, 70910-900, Brasilia D.F., Brazil; Graduate Program in Business Administration, University of Brasilia, 70910-900, Brasilia D.F., Brazil

**Keywords:** Birds, Rank abundance distribution, Species abundance distribution, Power-law distribution

## Abstract

Only a few bird species are abundant. According to new data, a log left-skewed distribution, rather than a lognormal distribution, better adjusts to the abundance distribution of bird species. We look at the rank abundance distribution rather than the species abundance distribution and find three power laws that fit the tails using the same data.

## Introduction

One of the few universal patterns in ecology is the distribution of species abundance (Morlon et al. 2009). The fact that some species are common but many more are rare is a crucial aspect of life’s diversity (Enquist et al. 2019). The species abundance distribution has a consistent general form, with many rare species coexisting with a few abundant species. The lognormal distribution is widely accepted in the ecology literature for the distribution of species abundance (Chisholm 2007). A lognormal distribution emerges when the number of individuals or another measure of abundance is plotted in a log scale, usually of base 2.

Callaghan et al. (2021) used the most recent influx of citizen science data to estimate the number of individuals (abundance) for 9,700 bird species, accounting for roughly 92 percent of all birds. They merged counts from the app eBird (https://ebird.org/home), which allows users to record bird sightings, with data from 724 well-studied species. They then utilized an algorithm to extrapolate sample estimates. As a result, they discovered a large number of species with small populations confined in niche habitats, as well as a limited number of species spread throughout a large area.

Callaghan et al. (2021) note that few of the species in their data are extremely abundant, and most have low population estimates. However, they also note that there would be about 200 more bird species than anticipated if the abundance of bird species followed a true lognormal distribution. They, therefore, propose a log left-skewed distribution.

Estimates of bird species abundance are critical for ecology, evolutionary biology, and conservation, and progress in quantifying abundance is welcome. Rather than the distribution of bird species abundance, we are interested in the distribution of bird rank abundance using the same data as Callaghan et al. (2021). We show that power laws fit the data well.

## Materials and methods

The data is depicted in Figure 1a. There are four undomesticated species with billions of individuals among the 50 billion birds: house sparrows (*Passer domesticus*), European starlings (*Sturnus vulgaris*), ring-billed gulls (*Larus delawarensis*), and barn swallows (*Hirundo rustica*). In comparison, each of the 5,022 species has fewer than 500,000 birds. Figure 1a shows a glimpse of the power-law pattern we discover and immediately draws attention to the group of 1,178 species that appear to deviate from the pattern. The abundance of a species is shaped by a variety of ecological processes (McGill et al. 2007). Many rarer species may have evolved to colonize a single island, for example, but human activity like deforestation can also explain the pattern deviation in Figure 1a. Some species may be rare as a result of our interference. From the standpoint of conservation initiatives, this issue needs to be looked into further.

**Figure 1.**
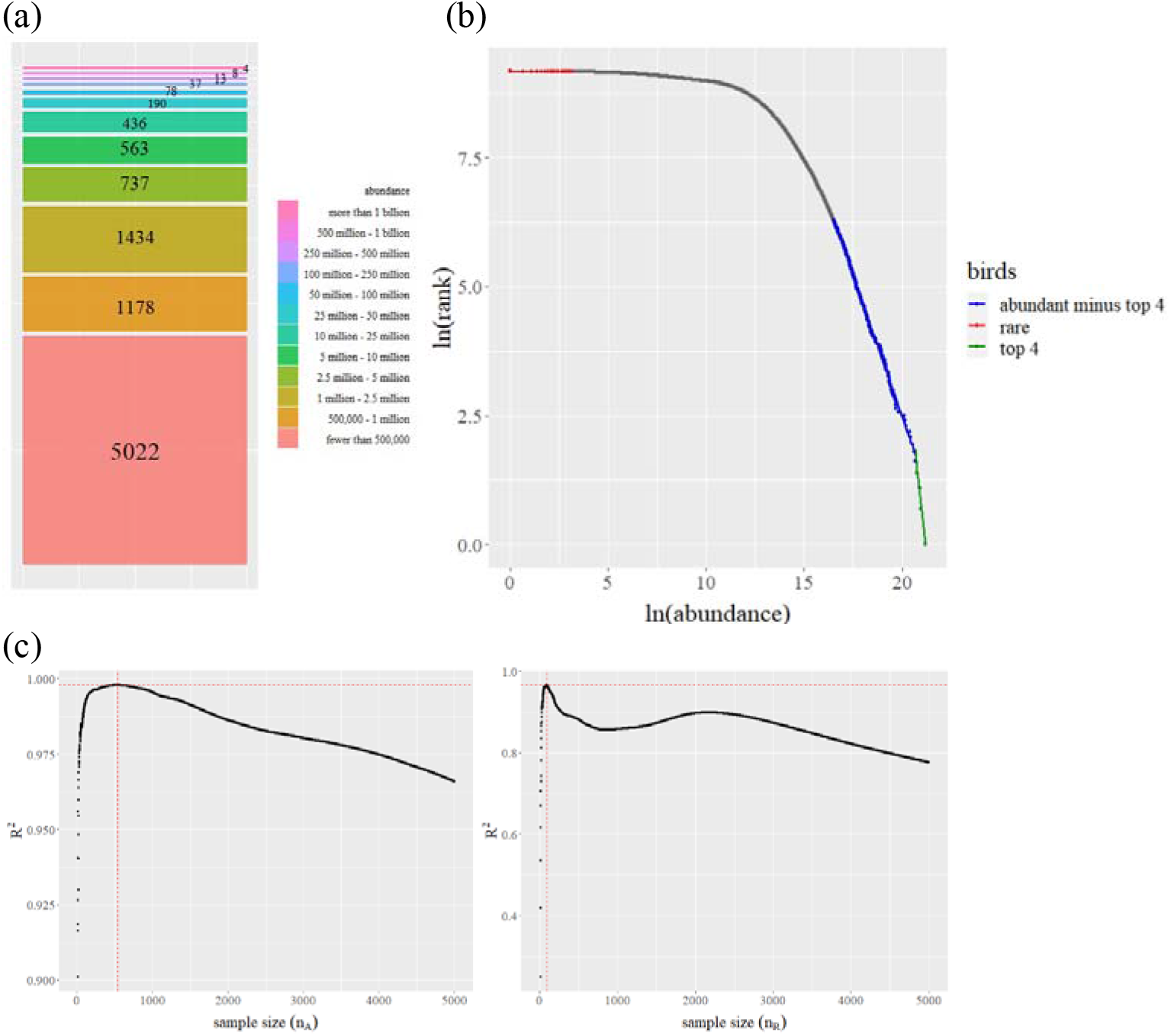
(a) Only a few bird species are abundant. (b) Three power laws emerge when we take ln(rank) versus ln(abundance). (c) *R*^2^ values versus sample size for the most abundant *n_A_* species (apart from the top four). The crossing of the dotted lines indicates the optimal sample size (*n_A_* = 534) that provides a maximum*R*^2^ = 0.998 (left). *R*^2^ values versus sample size for the rarer *n_R_* species; the optimal sample size is *n_R_* = 89 with a maximum *R*^2^ = 0.966 (right).

A power law governs a quantity when the probability of receiving a given value varies inversely with its power (Newman 2005). Thus, a power law is a relationship between two quantities in which a change in one causes a proportional change in the other. This holds true regardless of the starting values. In the dynamics of hierarchical systems that arise in physics, economics, and biology, power laws are common. A power law appears geometrically as a straight line in a plot of log of rank and log of the quantity at hand. After ranking the bird species from top to bottom, three power laws emerge. Figure 1b is the result of plotting the log of rank against the log of bird abundance. The power laws in Figure 1b can be quantified by calculating their Pareto exponents in relation to the straight line slopes.

One common approach is to use the survival function, *S*(*x*), in a Pareto type I model (Jenkins 2017). Those bird species whose abundance exceeds *x* – that is, one minus the cumulative distribution function *P*(*x*) – are given by

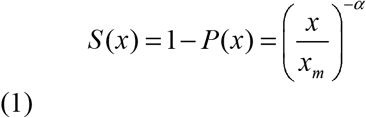

where *x_m_* is the lower bound on abundance, and *x* ≥ *x_m_* > 0. The Pareto exponent (tail index) is a shape parameter *α* that describes the heaviness of the right tail, with smaller values indicating a heavier tail.

## Results

We can estimate the three Pareto exponents associated with the rare and abundant species by ordinary least squares regressions of log of the survivor function on log of abundance and a constant term (Jenkins 2017). These result in straight-line intervals of slope −*α* (Figure 1b).

To determine the group of the most abundant *n_A_* species (aside from the top four), we run linear regressions varying *n_A_* from 10 to 5000 and record the corresponding values of the coefficient of determination *R*^2^. Then, we do the same for the *n_R_* rarer species, also ranging *n_R_* from 10 to 5000. Figure 1c shows the results. For the most abundant species, model (1) fits well for the *n_A_* = 534 most abundant species, providing *R*^2^ = 0.998. As for the rarest species, the best fit presents *R*^2^ = 0.966 for the *n_R_* = 89 rarest species.

Table 1 shows the Pareto exponents for the three subsamples. The value of *α* very close to zero for rare species suggests a heavier tail. Thus, these species are in a high uncertainty zone because there is no expected value for the number of rare species *X*. As a result, such species are in a non-equilibrium state in which they can be on the verge of extinction (*X* = 0) or becoming increasingly rare. Apart from the top four, the distribution of the most abundant species has an expected value *X* for abundance. The distribution, however, lacks a clear variability (as *α* < 2), implying that this group is still vulnerable to abrupt hierarchical rank adjustments. Finally, the top four have the shortest tails. This finding is consistent with the literature, which shows that the risk of extinction varies across the avian phylogeny (Purvis 2008).

**Table 1.** Finding the Pareto exponents

| Group | Number of species | $R^2$ | Pareto exponent $\alpha$ | $\ln x_m$ |
| --- | --- | --- | --- | --- |
| Top four | 4 | 0.972 | 3.485 ($\pm 0.422$) | 73.90 ($\pm 8.85$) |
| Abundant minus top four | 546 | 0.998 | 1.107 ($\pm 0.002$) | 24.62 ($\pm 0.04$) |
| Rare | 89 | 0.966 | 0.0027 ($\pm 0.00005$) | 9.18 ($\pm 0.0001$) |

Figure 2 depicts a hypothetical evolution of the values of the Pareto exponent for the top four species as the *k* most abundant species are gradually removed. Because no abundant species are removed in the benchmark of Figure 1b, *k* = 0. The impact on the value of the Pareto exponent of removing the most abundant species is represented by *k* = 1, and the impact on the remaining top four is shown in Figure 2. The Pareto exponents are then computed after removing the two most abundant species (*k* = 2), the three most abundant (*k* = 3), and so on. Figure 2 shows a Loess smooth curve fitting with a blue line. The shaded area represents a 95% confidence interval for *α*.

**Figure 2.**
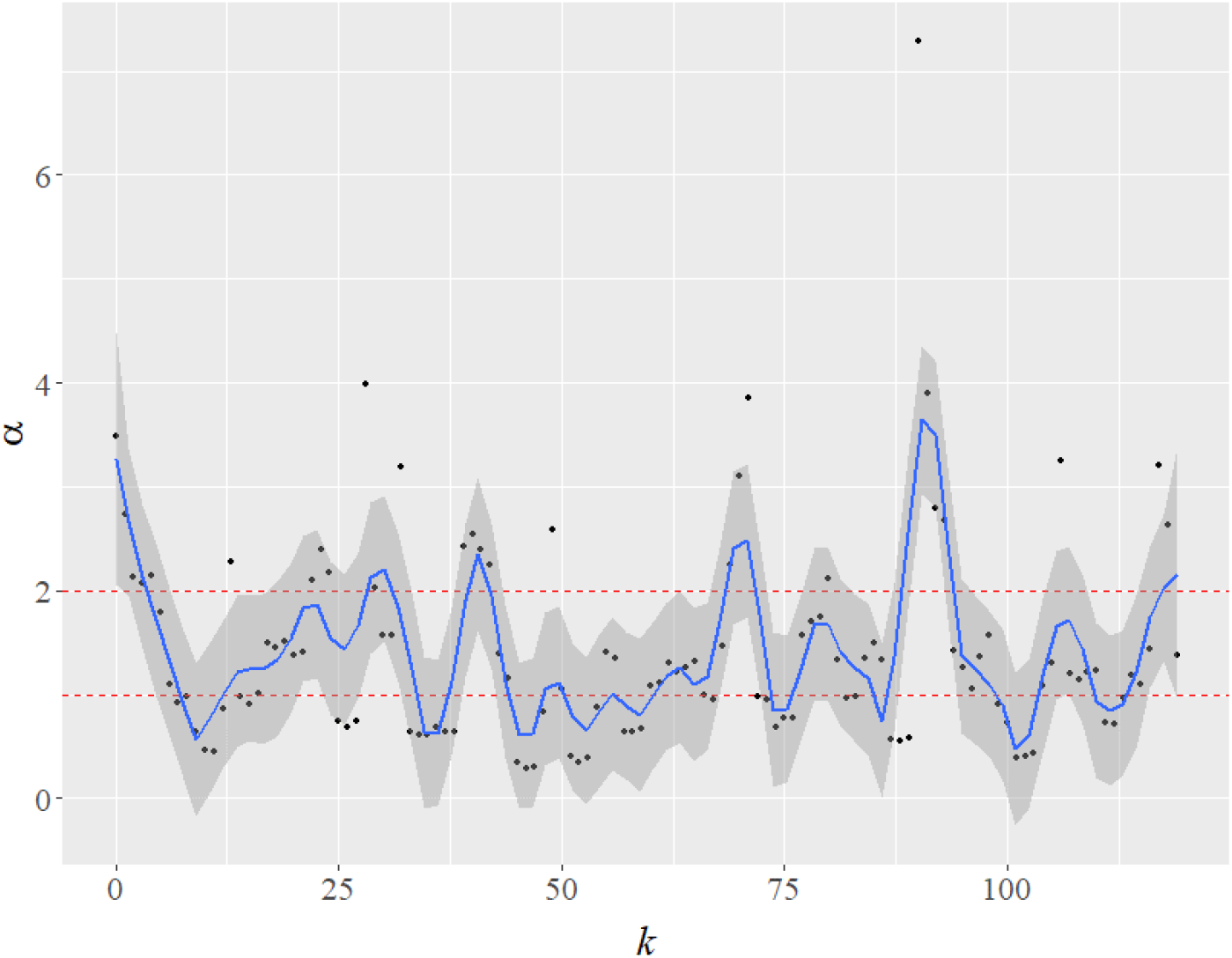
Changes in the value of the Pareto exponent (*α*) as we simulate the impact of the extinction of the most abundant species (as *k* increases) on the other remaining four large species.

The results of the exercise show a pattern of interest in the impact of extinction on the remaining (top four) species. The Pareto exponents, in particular, become unstable. The alphas are oscillating between light-tailed (*α* > 2) and heavy-tailed regimes. Of note, when 1 < *α* ≤ 2 the Pareto has no variance and when *α* ≤ 1 it has no mean either. As a result, the extinction of the big four significantly increases the overall uncertainty for the other species.

## Concluding remarks

Only a few bird species are abundant. Using new data, Callaghan et al. (2021) propose that a log left-skewed distribution, rather than a lognormal distribution, adjusts better to the abundance distribution of bird species. Using the same data, we consider the rank abundance distribution rather than the species abundance distribution. Three power laws are found to fit the tails. Because power laws emerge even when data is scarce, our finding is not overly dependent on data quality.

The fact that many species, like the third-most abundant ring-billed gull, have a Nearctic distribution among the top ten most abundant species may be one limitation of the Callaghan et al.’s data. Because of this supposed limitation, the eBird database could not reliably estimate the abundance of most species that do not occur widely in the Americas. However, this criticism ignores the expert ornithologist knowledge that went into compiling the dataset, and we cannot know any better.

## Notes

**Funding**. This work was supported by CNPq, Capes, FAP-DF, and DPI-DPG UnB.

https://www.pnas.org/highwire/filestream/984963/field_highwire_adjunct_files/1/pnas.2023170118.sd01.xlsx

### Competing Interest Statement

The authors have declared no competing interest.

### Summary of Updates

The presentation and concepts have been clarified in the revised manuscript.

